# LIX1 controls digestive mesenchyme-derived cell fate decision by regulating cristae organization in mitochondria

**DOI:** 10.1101/2022.01.06.475258

**Authors:** Amandine Guérin, Claire Angebault, Sandrina Kinet, Chantal Cazevieille, Manuel Rojo, Jérémy Fauconnier, Alain Lacampagne, Arnaud Mourier, Naomi Taylor, Pascal de Santa Barbara, Sandrine Faure

## Abstract

Limb Expression 1 (LIX1) is a master regulator of digestive mesenchymal progenitor and GastroIntestinal Stromal Tumor (GIST) cell proliferation by controlling the expression of the Hippo effectors YAP1/TAZ and KIT. However, the underlying mechanisms of these LIX1-mediated regulations and tumor promotion remain to be elucidated. Here, we report that LIX1 is S-palmitoylated on cysteine 84 and localized in mitochondria. LIX1 knock-down affects the mitochondrial ultrastructure, resulting in decreased respiration and mitochondrial reactive oxygen species production. This is sufficient to downregulate YAP1/TAZ and reprogram KIT-positive GIST cells towards the smooth muscle cell lineage with reduced proliferative and invasive capacities. Mechanistically, LIX1 knock-down impairs the stability of the mitochondrial proteins PHB2 and OPA1 that are found in complexes with mitochondrial-specific phospholipids and are required for cristae organization. Supplementation with unsaturated fatty acids counteracts the effects of LIX1 knock-down on mitochondrial morphology and ultrastructure, restores YAP1/TAZ signaling, and consequently KIT levels. Altogether, our findings demonstrate that LIX1 contributes to GIST aggressive potential by modulating YAP1/TAZ and KIT levels, a process that depends on mitochondrial remodeling. Our work brings new insights into the mechanisms that could be targeted in tumors in which YAP1 and TAZ are implicated.

## 1. Introduction

GISTs originate from Interstitial Cells of Cajal (ICCs) that share progenitors with smooth muscle cells (SMCs) [1,2]. GISTs are the most common mesenchymal neoplasm in the gastrointestinal tract and represent ∼5% of all sarcomas [3,4]. GISTs occur predominantly in the stomach (50–60%) and small intestine (30–35%), and result from the deregulated proliferation of ICCs [5]. More than 80% of GISTs harbor a gain of function mutation in *KIT*, and less frequently in Platelet-Derived Growth Factor Receptor alpha (*PDGFRA*). These mutations result in the constitutive activation of these tyrosine kinase receptors and their signaling pathways, leading to spontaneous proliferation and uncontrolled tumor growth [6]. For patients with metastatic or unresectable GIST, molecular targeted therapies based on tyrosine kinase inhibitors, imatinib mesylate (Gleevec®) is the only efficient treatment [7]. However, the majority of patients develop resistance to imatinib within 2-3 years [8]. Therefore, the molecular mechanisms driving digestive mesenchymal cell differentiation and plasticity must be identified to understand GIST development.

Cell dedifferentiation is a central mechanism in the initiation of neoplastic transformation [9–11]. It involves loss of lineage-specific gene expression and regression from a specialized function to a more primitive state through expression of genes that govern embryonic cell-fate specification. Such mechanism is also observed in Gastrointestinal stromal tumors (GISTs), as demonstrated by the high expression of LImb eXpression 1 (*LIX1)*, a mesenchymal progenitor marker that controls mesenchyme-derived cell fate decision within the digestive musculature. Therefore, LIX1 is an attractive novel target for GIST therapeutics [12,13].

The human *LIX1* gene encodes a 282-amino acid protein. LIX1 was discovered in chick embryos in a screen to identify markers of limb development [14]. An *in silico* prediction analysis indicated that LIX1 has a double-stranded RNA binding domain, suggesting that it could be involved in RNA processing [15]. Moreover, the arthropod homolog of *LIX1, lowfat*, interacts with the atypical cadherins *fat* and *dachsous*, two upstream components of the Hippo pathway implicated in maintaining tissue homeostasis through the regulation of the balance between cell proliferation and differentiation [15,16]. Accumulating evidences indicate that LIX1 is a key regulator of muscle progenitor proliferation in vertebrates [12,17]. In the digestive musculature, LIX1 is only expressed during fetal life. In this tissue, it stimulates the expression and activity of the Hippo effector YAP1, and LIX1 and YAP1 are key regulators of stomach mesenchymal progenitor development [12]. Moreover, LIX1 expression is high in GISTs and is associated with poor prognosis. *LIX1* silencing in GIST cells in culture and in a xenograft model decreases YAP1/TAZ expression and activity, reduces the expression of KIT and of genes involved in GIST aggressiveness, and reprograms KIT-positive GIST cells to the SMC lineage, thus decreasing tumor cell proliferation and invasion [13]. However, the exact mechanisms by which LIX1 contributes to YAP1/TAZ regulation remain to be elucidated.

Mitochondria are signaling organelles that regulate many cellular events and can dictate cell fate [18]. Mitochondria have many cellular functions, including coupling respiration to ATP synthesis *via* oxidative phosphorylation (OXPHOS), an activity associated with ROS production that is implicated in the regulation of epigenetic signaling and cell growth control [19,20]. Accumulating evidences indicate that mitochondria orchestrate the metabolic rewiring required for neoplastic growth [21].

Here, we found that LIX1 is a S-palmitoylated protein with mitochondrial localization. LIX1 knock-down in GIST cells affected mitochondrial morphology and ultrastructure, resulting in decreased respiration and mitochondrial reactive oxygen species (mtROS) production, two key mitochondrial functions involved in cancer cell proliferation and tumorigenesis. Our findings demonstrated that LIX1 regulates YAP1/TAZ levels and plasticity between the ICC and SMC lineages by controlling mitochondrial functions.

## 2. Results

### 2.1. LIX1 is tightly anchored to the outer membrane of mitochondria

To better understand LIX1 role in GIST, we first analyzed its localization by confocal microscopy in HeLa cells that express HA-LIX1. We found that a significant fraction of HA-LIX1 co-localized with the mitochondrial marker COXIV (Fig. 1A,B). We then performed fractionation experiments using control non-transfected cells (empty) and HA-LIX1-expressing cells to analyze LIX1 expression in the total protein extract, cytoplasmic (crude cyto) and mitochondrial (mito) fractions. Western blot analysis with antibodies against GAPDH (cytosol marker), TFAM (mitochondrial marker) and LIX1 showed that LIX1 was a cytoplasmic protein enriched in the mitochondrial fraction (Fig. 1C). Mitochondria include an outer mitochondrial membrane (OMM) that surrounds the inner mitochondrial membrane (IMM), with a small intermembrane space (IMS) in between. The IMM forms many folds (cristae) that protrude into the internal compartment (or matrix). We next determined LIX1 sub-mitochondrial localization using a PK accessibility test and alkali treatments. Incubation of isolated mitochondria from HA-LIX1-expressing HeLa cells with increasing PK concentrations did not affect the levels of proteins localized in the IMS (CYT C), IMM (COXIV), and matrix (TFAM). Conversely, it strongly reduced VDAC and TOM20, two OMM proteins, and also LIX1 levels (Fig. 1D). This suggested that LIX1 is localized at the OMM surface. To determine the strength of LIX1 interaction with the OMM, mitochondrial fractions from HA-LIX1-expressing cells were incubated with sodium carbonate (pH11.5). This treatment at high pH disrupts protein-protein and weak protein-lipid interactions, and leads to membrane integrity loss. As expected, we retrieved CYT C, an IMS protein, in the soluble (S) faction, whereas transmembrane proteins, such as COXIV, remained in the pellet (P)/membrane fraction as well as HA-LIX1 (Fig. 1E). Furthermore, incubation of mitochondria purified from HA-LIX1-expressing cells with nonionic detergents (digitonin, NP-40, Triton X-100) did not affect LIX1 level, even at concentrations that extract partially or totally other mitochondrial proteins, such as TFAM (matrix), TOM20 (OMM) and COXIV (IMM) (Fig. 1F). Indeed, we detected mitochondrial LIX1 in the Pellet fraction as well as VDAC, a protein localized at mitochondrial detergent-resistant micro-domains. Both VDAC and LIX1 were only partially dissociated, even upon incubation with 1% SDS, a concentration that fully dissociated TOM20 and COXIV from the OMM (Fig. 1F). This indicated that LIX1 is strongly attached to the OMM, presumably in detergent-resistant micro-domains. These raft-like micro-domains, enriched in the phospholipid cardiolipin, are located at the contact sites between the IMM and OMM [22] and play important roles in mitochondrial metabolism regulation.

**Figure 1.**
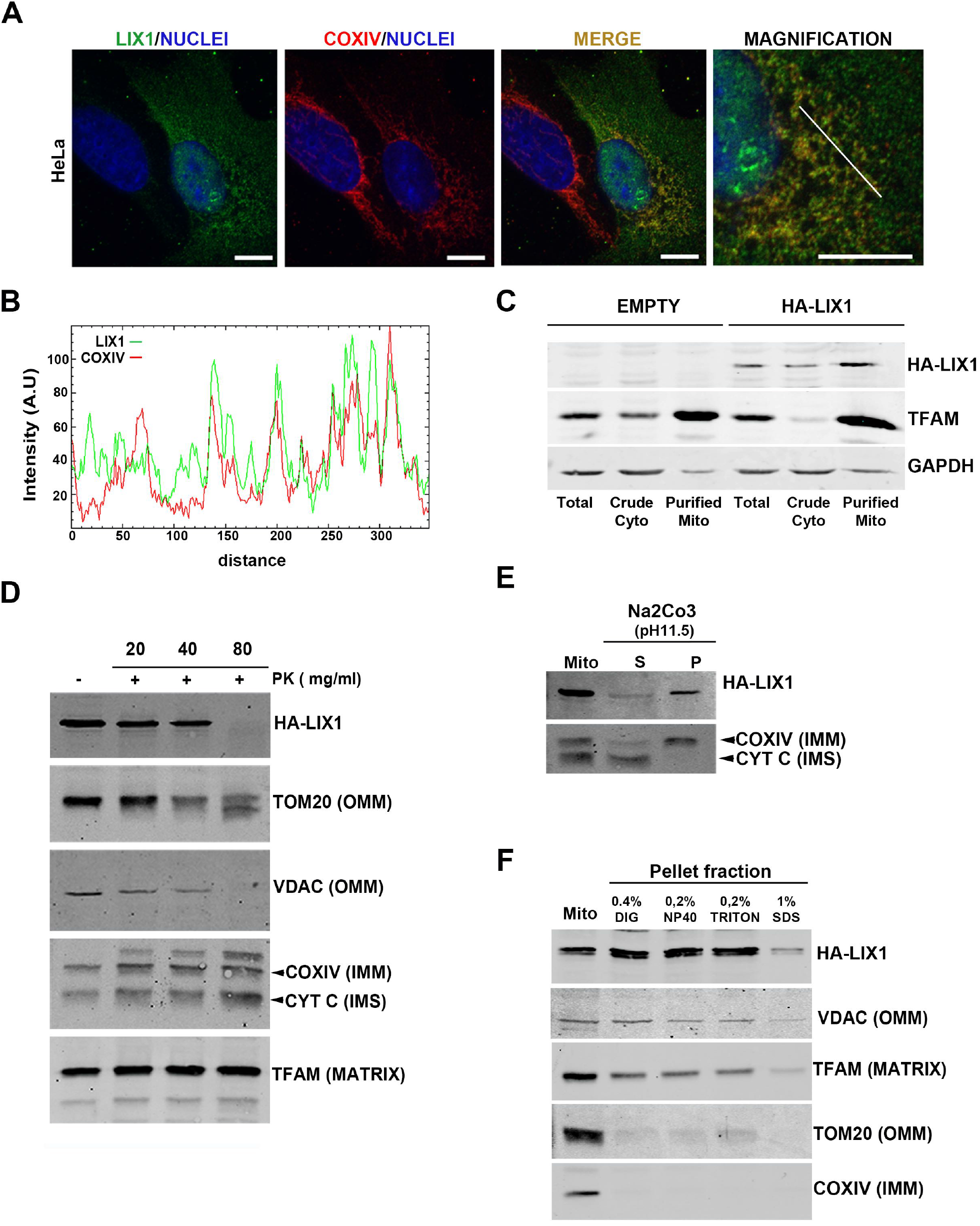
LIX1 localizes to mitochondria, and is tightly anchored to the outer membrane. (**A**) Confocal microscopy analysis of HA-LIX1 localization in HeLa cells. Comparison with the mitochondrial marker COXIV. Scale bars, 10μm. (**B**) RGB profile of LIX1 and COXIV expression co-localization. (**C**) Western blot analysis of total protein extracts (Total), crude cytoplasmic (Crude Cyto) and purified mitochondria (Purified Mito) from non-transfected HeLa cells (Empty) and HeLa cells expressing HA-LIX1. Membranes were probed with antibodies against the HA-tag, GAPDH (cytoplasmic) and TFAM (mitochondrial marker). (**D**) Sub-mitochondrial localization of HA-LIX1 in isolated mitochondria from HA-LIX1-expressing HeLa cells after (+) or not (-) proteinase K (PK) accessibility test. Membranes were probed with antibodies against the HA-tag, TOM20 and VDAC (Outer Mitochondrial Membrane (OMM) markers), COXIV (Inner Mitochondrial Membrane (IMM) marker), Cytochrome C (CYT C) (Inter-Membrane Space (IMS) marker) and TFAM (matrix protein). (**E**) Western blot analysis of total isolated mitochondria (Mito) from HA-LIX1-expressing cells after incubation with Na_2_CO_3_ pH11.5, and ultracentrifugation to separate the supernatant (S) and pellet (P) fractions. Membranes were probed with antibodies against COXIV (IMM marker) and cytochrome C (CYT C) (IMS soluble marker). (**F**) Western blot analysis of total isolated mitochondria (Mito) from HA-LIX1-expressing HeLa cells after solubilization (+) or not (-) with the indicated detergents and ultracentrifugation. Membranes were probed with antibodies against the HA-tag, TFAM (matrix protein), VDAC and TOM20 (OMM markers) and COXIV (IMM marker).

### 2.2. LIX1 mitochondrial localization is controlled by S-palmitoylation on cysteine 84

As LIX1 does not have any transmembrane domain, we next wanted to determine how LIX1 association with the OMM is regulated. Accumulating evidences indicate that lipid modifications can mediate protein translocation to the OMM. For instance, S-palmitoylation is required for BAX targeting to the OMM to initiate apoptosis [23]. The GPS-Lipid algorithm (http://lipid.biocuckoo.org), which allows predicting four lipid modifications (S-palmitoylation, N-myristoylation, S-farnesylation and S-geranylgeranylation) [24], identified two potential S-palmitoylation sites (cysteine 83 and 84) in LIX1 (Fig. 2A). S-palmitoylation is a dynamically regulated post-translational modification in which palmitic acid (C16.0) is reversibly attached to a cysteine residue *via* a thioester bond [25]. Upon S-palmitoylation, cytosolic proteins acquire a hydrophobic anchor that facilitates their docking to membranes, presumably into raft-like micro-domains [26]. To determine whether LIX1 is S-palmitoylated, we transfected HeLa cells with wild-type HA-LIX1 (LIX1 WT) or LIX1 mutants in which cysteine 83 and 84 were replaced by serine residues (LIX1 C83S, LIX1 C84S). We then tested S-palmitoylation in total protein extracts using the acyl-RAC assay [27,28]. A specific signal corresponding to S-palmitoylation was detected in the cleaved bound fraction (cBF) of LIX1 WT and LIX1 C83S samples, but not in the LIX1 C84S sample (Fig. 2B). Next, to determine whether cysteine 84 is required for LIX1 translocation to mitochondria, we analyzed by western blotting total extracts and purified mitochondrial fractions of HeLa cells that express WT HA-LIX1, LIX1 C83S, LIX1 C84S, or the double mutant LIX1 C83S/C84S. Mitochondrial targeting was strongly reduced (by ∼50%) for LIX1 C84S and LIX1 C83S/C84S compared to LIX1 WT and LIX1 C83S (Fig. 2C, D). These data indicate that LIX1 S-palmitoylation on cysteine 84 is implicated in its translocation to the OMM. The 3D structure of the LIX1 protein, established using the AlphaFold algorithm (https://alphafold.ebi.ac.uk/entry/Q8N485), revealed that the cysteine residue 84, located between one α-helix and one β-strand, is accessible for lipid modification (Fig. 2E). This cysteine residue is conserved in all vertebrate LIX1 orthologs and in the *Drosophila* homologs, but not cysteine 83 (Fig. 2A; Fig. S1).

**Figure 2.**
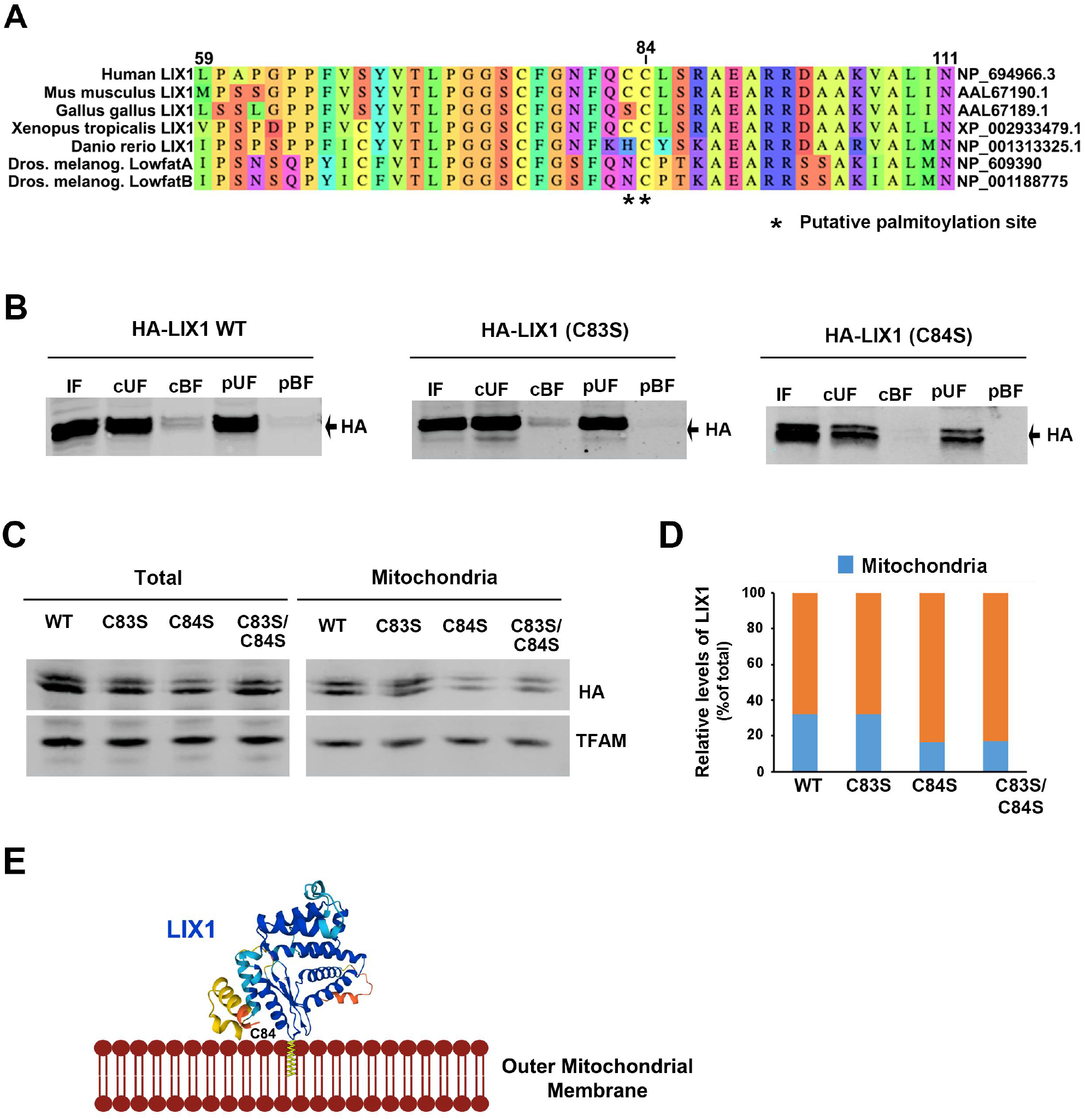
LIX1 mitochondrial localization is controlled by S-palmitoylation at Cys84. (**A**) Sequence alignment of human LIX1 and its orthologs. Asterisks indicate the position of the cysteine residues corresponding to putative S-palmitoylation sites. (**B**) LIX1 S-palmitoylation analysis with the Acyl RAC method. Membranes were incubated with anti-HA-tag antibodies to detect LIX1. IF, input fraction; cUF, cleaved unbound fraction; cBF, cleaved bound fraction; pUF, preserved unbound fraction; pBF, preserved bound fraction. **(C**) Western blot analysis of total extracts or isolated mitochondria from the same amount of total extract from HeLa cells that express WT HA-LIX1 (LIX1 WT), single LIX1 mutants (LIX1 C83S, LIX1 C84S) or the double mutant (LIX1 C83S/C84S) with antibodies against the HA-tag and TFAM (matrix protein). **(D**) Quantification of the western blots showing the percentage of LIX1 (normalized to TFAM level) at mitochondria relative to LIX1 level in the total extract (normalized to TFAM level). **(E**) 3D structure of the LIX1 protein showing that the cysteine residue 84 is accessible for modification.

As ectopically expressed LIX1 localized to mitochondria and as accumulating evidences points to mitochondria to orchestrate the metabolic rewiring required for neoplastic growth [21], next we investigated endogenous LIX1 expression. As we previously showed, LIX1 is highly expressed in gastric mesenchymal (gMes) progenitors and GIST-T1 cells as well as in the non-cancer cell line HEK-293T, but not in HeLa cells (Fig. 3A) [13]. Most importantly, analysis of LIX1 expression revealed that endogenous LIX1 is enriched in the mitochondrial fraction of GIST-T1 cells and gMes progenitor cells (Fig. 3B-D). LIX1 expression in mitochondria was strongly reduced in GIST-T1 cells that stably express *Sh*RNAs against *LIX1* (GIST-T1-*ShLIX1#2*) (Fig. 3D)[13]. Confocal analysis of GIST-T1 cells using the mitochondrial markers TOM20 and COXIV confirmed the mitochondrial localization of a significant fraction of endogenous LIX1 (Fig. 3E-H). Our data demonstrate that endogenous LIX1 is enriched in mitochondria in digestive mesenchymal progenitors and their mesenchymal-derived cells.

**Figure 3.**
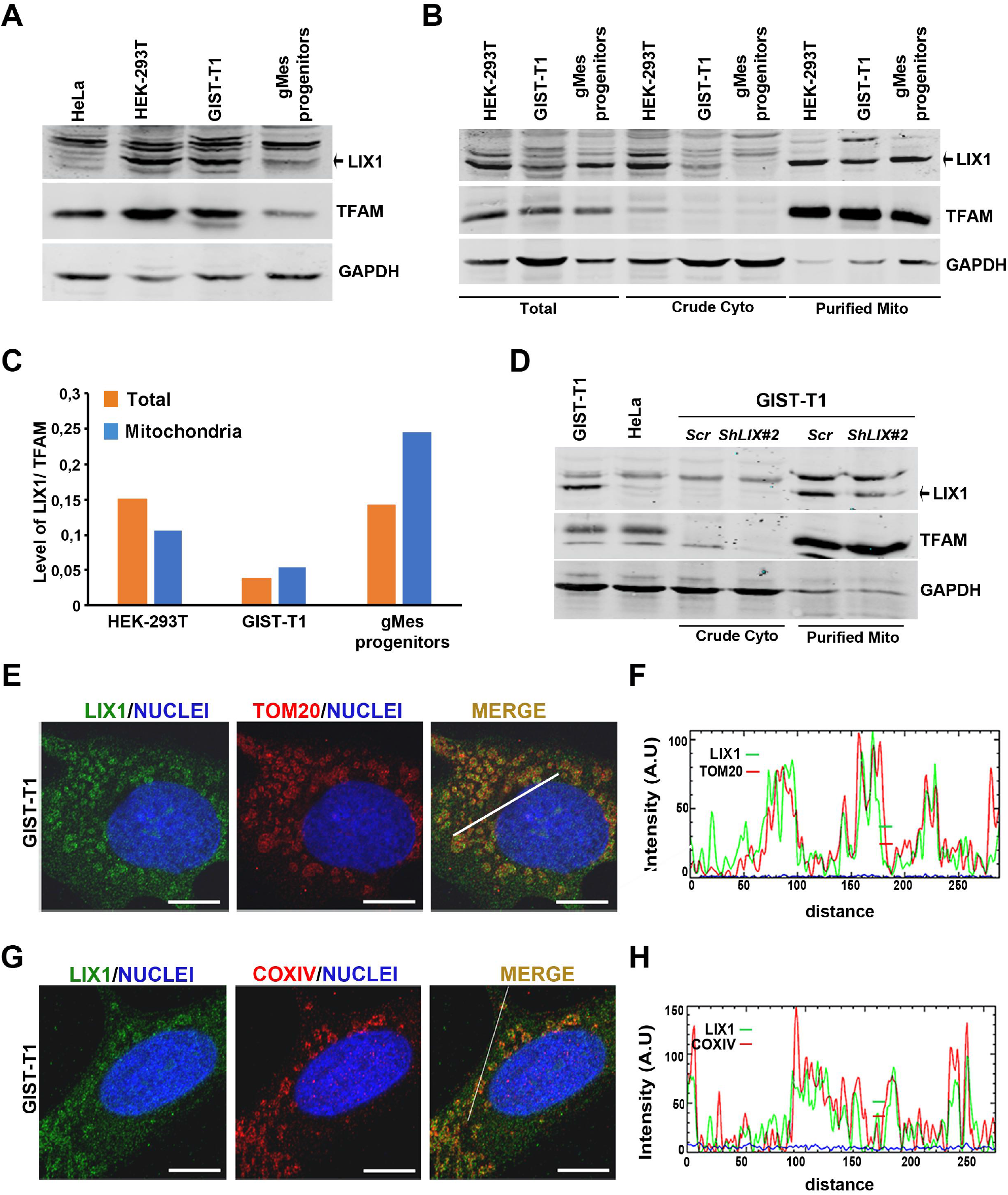
Endogenous LIX1 is enriched in mitochondria of GIST-T1 cells. (**A**) Western blot analysis of total extracts from HeLa, HEK-293T, GIST-T1, and gastric mesenchymal progenitor cells (gMes progenitors). Membranes were probed with antibodies against LIX1 and TFAM (mitochondrial marker). Equal loading was verified by GAPDH expression. (**B**) Western blot analysis of total extracts, crude cytoplasmic (Crude Cyto) and purified mitochondria (Purified Mito) fractions (equal amounts) from HEK-293T, GIST-T1 and gMes progenitors. Membranes were probed with antibodies against LIX1, GAPDH (cytoplasmic) and TFAM (mitochondrial marker). **(C**) Quantification in the western blot images of endogenous LIX1 (normalized to TFAM level) in total extracts and purified mitochondria. (**D**) Western blot analysis of total extracts from GIST-T1 and HeLa (negative control) cells, and of cytoplasmic (Crude Cyto) and mitochondrial (Purified Mito) fractions from GIST-T1 cells that stably express shRNAs against *LIX1* (*ShLIX1#2*) or scramble shRNAs (Scr). Membranes were probed with antibodies against LIX1, GAPDH (cytoplasmic) and TFAM (mitochondrial marker). (**E**) Confocal microscopy analysis of endogenous LIX1 localization in GIST-T1 cells. Comparison with the mitochondrial marker TOM20. (**F**) LIX1/TOM20 RGB profile analyzed at the level of the white line. (**G**) Confocal microscopy analysis of endogenous LIX1 localization in GIST-T1 cells. Comparison with the mitochondrial marker COXIV. (**H**) LIX1/COXIV RGB profile analyzed at the level of the white line.

### 2.3. LIX1 regulates mitochondrial respiration and ROS production in GIST cells

The mitochondrial localization of LIX1 in GIST-T1 cells prompted us to investigate its role in mitochondrial function. Therefore, we assessed LIX1 silencing effect on Oxygen consumption rate (OCR) and respiratory chain complex activities by measuring mitochondrial respiration. Extracellular flux (Seahorse) analysis showed that basal OCR (baseline minus non-mitochondrial respiration) and ATP-linked OCR (difference between OCR at baseline and respiration following oligomycin addition) were significantly lower in both GIST-T1-*ShLIX1* cell lines than in control GIST-T1-*Scrambled* cells (Fig. 4A, B). This indicated lower reliance on OXPHOS for energy production in GIST cells in which LIX1 was silenced. OCR quantification in permeabilized GIST-T1 cells demonstrated a significantly lower CXI-driven oxygen consumption in both GIST-T1-*ShLIX1* cell lines (Fig. 4D). Conversely, CXII-driven oxygen consumption was moderately decreased only in GIST-T1-*ShLIX1#2* but not in GIST-T1-*ShLIX1#1* cells, possibly due to the better silencing efficiency of *ShLIX1#2* [13]. The decreased OCR could not be explained by changes in OXPHOS protein expression as the expression levels of several respiratory chain subunits was comparable in silenced and control cells (Fig. 4C). Further, mitochondrial ROS (mtROS) production, measured by immunofluorescence analysis with the MitoSOX probe, was decreased in GIST-T1-*ShLIX1* cells compared with control (Fig. 4E, F). These data indicate that reducing LIX1 levels mainly interferes with CXI-driven respiration, leading to decreased mtROS production.

**Figure 4.**
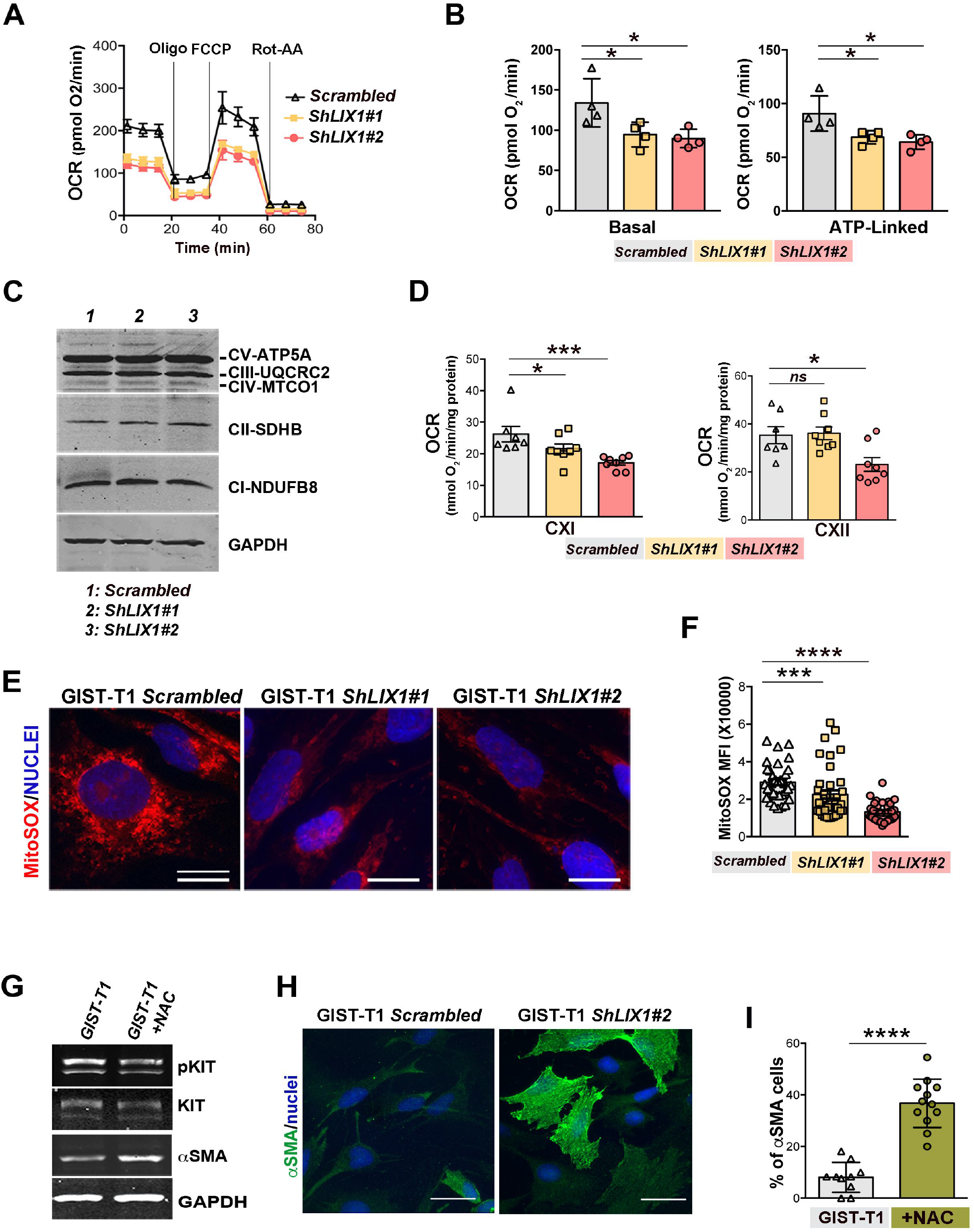
LIX1 regulates mitochondrial respiration and ROS production in GIST-T1 cells. (**A**) Oxygen consumption rate (OCR) in GIST-T1-*Scrambled*, GIST-T1-*ShLIX1#1* and GIST-T1-*ShLIX1#2* cells incubated with oligomycin (Oligo), FCCP and rotenone/antimycin A (Rot-AA) using a Seahorse XF96 Extracellular Flux analyzer. Data are the mean ± SEM of 4 independent experiments. Cells were left for 4-6h in the plates before starting the injection of drugs. (**B**) Quantification of the basal and ATP-linked OCR values from the Seahorse analysis. Non-mitochondrial OCR was determined after the addition of rotenone/antimycin A (Rot-AA) and subtracted from all the other values before calculating the respiratory parameters. Values are the mean ± SD of *n=4* experiments using GIST-T1-*Scrambled*, GIST-T1-*ShLIX1#1* and - *ShLIX1#2* cells. *P <0.05 (two-tailed Mann–Whitney tests). (**C**) Western blot analysis of total protein extracts from GIST-T1-*Scrambled*, GIST-T1-*ShLIX1#1* and GIST-T1-*ShLIX1#2* cells. Membranes were probed with antibodies against the indicated OXPHOS proteins. (**D**) ETC-CXI- and ETC-CXII-driven respiration measured using Oxygraph-2k high-resolution respirometry. Values are the mean ± SEM of GIST-T1-*Scrambled* (*n=7)*, GIST-T1-*ShLIX1#1 (n=8)* and GIST-T1-*ShLIX1#2 (n=8)* cells. *P <0.05, ***P < 0.001 (one-tailed Mann–Whitney test). (**E**) MitoSOX staining of GIST-T1-*Scrambled*, GIST-T1-*ShLIX1#1* and -*ShLIX1#2* cells. Nuclei were visualized with Hoechst. Scale bars, 20μm. (**F**) Quantification of the MitoSOX signal. Values are the mean ± SEM of GIST-T1-*Scrambled (n=34)*, GIST-T1-*ShLIX1#1 (n=39)*, and GIST-T1-*ShLIX1#2 (n=35)* cells from three independent experiments. ***P <0.001; ****P <0.0001 (two-tailed Mann–Whitney test). (**G**) Representative western blot showing phosphorylated KIT(pKIT), KIT and αSMA levels in GIST-T1 cells incubated or not (control) with 3mM N-acetyl cysteine (NAC) for 18h. Equal loading was verified by GAPDH expression. (**H**) Immunofluorescence analysis of GIST-T1 and GIST-T1+NAC - cells with anti-αSMA antibodies. Nuclei were visualized with Hoechst. Scale bar, 50μm. (**I**) Quantification of the percentage of αSMA-positive cells. Values are the mean ± SEM of *n =*136 GIST-T1-*Scrambled* and *n=*125 for GIST-T1+NAC. ****P <0.0001 (two-tailed Mann–Whitney test).

High levels of mtROS activate signaling pathways that promote cancer cell proliferation [29–32]. Our previous findings indicated that LIX1 silencing promotes a phenotypic switch of GIST cells that express KIT (required for mesenchymal progenitor specification toward ICCs and GIST development) towards the SMC lineage, and reduces their tumorigenic and malignant potential [13]. Therefore, we hypothesized that LIX1 modulation of GIST cell identity could be mediated through its capacity to control mtROS production. In line with this hypothesis, and as observed in GIST-T1-*ShLIX1* cells, reducing mtROS level in GIST-T1 cells using N-acetyl cysteine (NAC), a ROS scavenger, led to a marked decrease of KIT expression coupled to the induction of the smooth muscle marker αSMA (Fig. 4G-I). These data highlight the pivotal role of the mitochondrial redox state regulation in the control of ICC specification and GIST tumorigenesis, a process regulated by LIX1.

### 2.4. LIX1 regulates the mitochondrial network morphology and the inner membrane architecture in GIST cells

As mitochondrial bioenergetics is influenced by the mitochondrial morphology and ultrastructure [33], we evaluated LIX1 implication in the regulation of mitochondrial morphology [34]. Under MitoTracker Red staining, observed fragmented mitochondria in GIST-T1-*Scrambled* cells, and an elongated network in GIST-T1-*ShLIX1* cells (Fig. 5A). Quantification of these findings showed that the number of mitochondria was reduced in LIX1-silenced compared with control cells (Fig. 5B). Again, the mitochondrial mass did not appear to be affected by LIX1 silencing (Fig. 4C). The decreased number of mitochondria in GIST-T1-*ShLIX1* cells was associated with a slight increase in the mean size, elongation and interconnectivity of mitochondria, compared with control cells (Fig. 5B). Conversely, HA-LIX1 expression in HeLa cells induced mitochondrial fragmentation, a phenotype that led to a significant increase in mitochondrial number and a decrease in mitochondrial elongation and interconnectivity (Fig. S2A-B). As the electron transport chain is in the IMM, we next analyzed the IMM architecture by TEM. Compared with GIST-T1-*Scrambled* cells, the IMM and cristae were altered in GIST-T1-*ShLIX1* cells (Fig. 5C; Fig. S3). These morphological changes were more important in GIST-T1-*ShLIX1#2* cells in which *LIX1* downregulation was more efficient. Specifically, lamellar cristae were almost completely lost and replaced by vesicle-like cristae without cristae junctions. We confirmed the remodeling of cristae in GIST-T1-*ShLIX1#2* cells by three-dimensional reconstructions from serial TEM images (Fig. 5D; Supplemental Movie 1 and Movie 2). Such aberrant balloon-like cristae were previously described in Prohibitin 2 (PHB2)-deficient mitochondria, where this phenotype was caused by the loss of the long OPA1 isoform, a dynamin-like GTPase required for mitochondrial fusion and cristae morphogenesis [35–37]. Analysis of PHB1 and PHB2 expression in total protein extracts and purified mitochondria from GIST-T1-*Scrambled* and GIST-T1-*ShLIX1* cells showed that their expression in total extracts was not affected by LIX1 silencing, whereas only PHB2 was decreased in mitochondria of GIST-T1-*ShLIX1* cells (Fig. 5E). This was associated with a reduction of all OPA1 isoforms in mitochondria (Fig. 5F). Our data demonstrate that LIX1 silencing in GIST-T1 cells impairs the stability of proteins required for proper cristae morphology and alters the mitochondrial membrane ultrastructure, resulting in decreased respiration and mtROS production.

**Figure 5.**
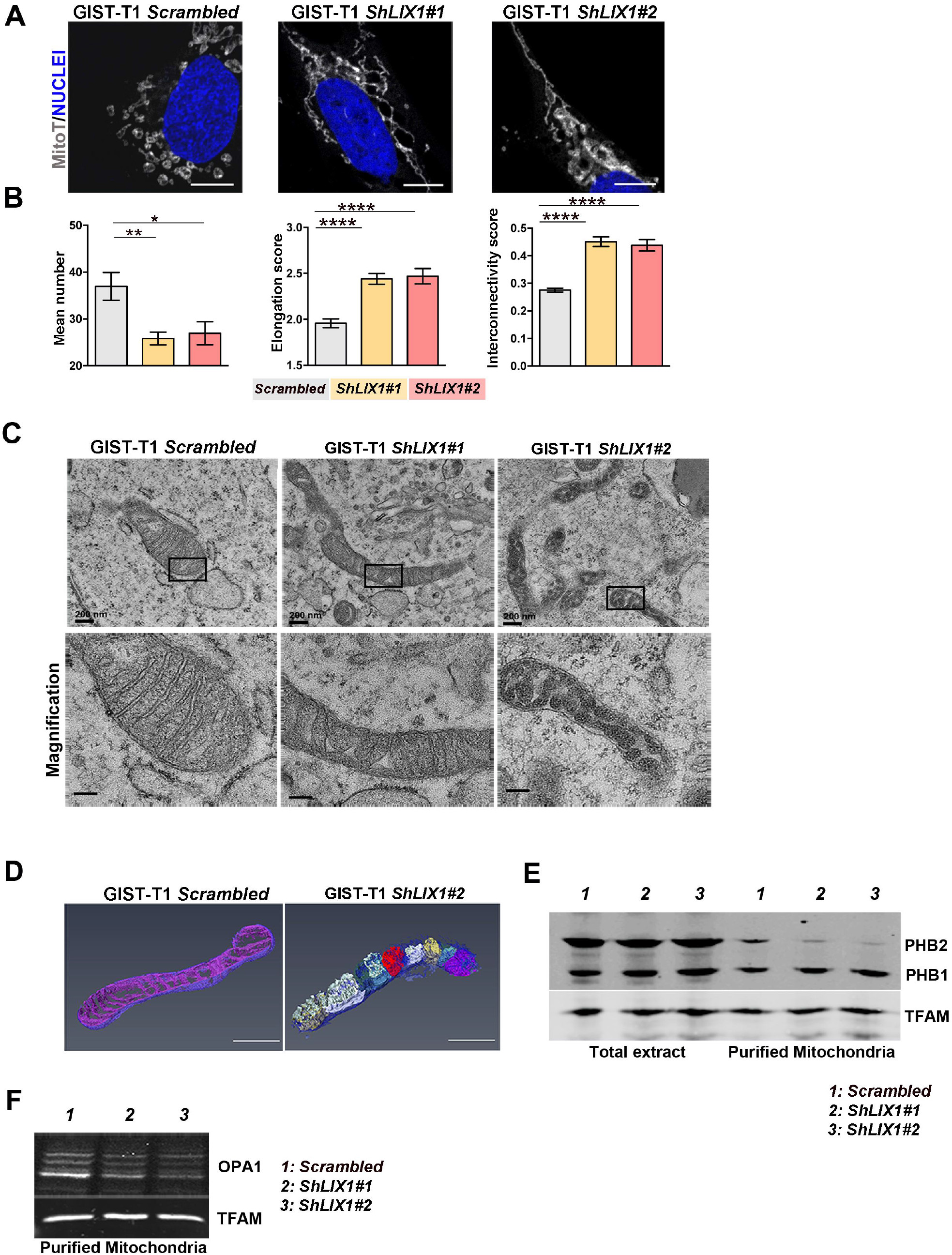
LIX1 regulates mitochondrial network morphology and inner membrane architecture in GIST-T1 cells. (**A**) MitoTracker staining of GIST-T1-*Scrambled*, GIST-T1-*ShLIX1#1* and -*ShLIX1#2* cells. Nuclei were visualized with Hoechst. Scale bars, 10μm. (**B**) Quantification of the MitoTracker data with the Mito-Morphology Macro in Image J. Values are the mean ± SEM of GIST-T1-*Scrambled (n=25)*, GIST-T1-*ShLIX1#1 (n=28)*, and GIST-T1-*ShLIX1#2 (n=28)* cells. *P <0.05; **P <0.001; ****P <0.0001 (two-tailed Mann–Whitney test). (**C**) Ultrastructure of mitochondria in GIST-T1-*Scrambled*, GIST-T1-*ShLIX1#1*, and - *ShLIX1#2* cells. Scale bars, 100 nm. Lower panels: magnifications of the areas in the black boxes in the upper panels. (**D**) Three-dimensional reconstructions from serial transmission electron microscopy sections of GIST-T1-*Scrambled* and GIST-T1-*ShLIX1#2* cells. Scale bars, 500nm. The vesicle-like cristae are shown in different colors. Not really clear what you want to show. (**E**) Western blot analysis of total protein extracts and purified mitochondria from GIST-T1-*Scrambled*, GIST-T1-*ShLIX1#1*, and -*ShLIX1#2* cells. Membranes were probed with anti-PHB and -TFAM (mitochondrial marker) antibodies. (**F**) Western blot analysis of total protein extracts from GIST-T1-*Scrambled*, GIST-T1-*ShLIX1#1*, and -*ShLIX1#2* cells. Membranes were probed with anti-OPA1 and -TFAM (mitochondrial marker) antibodies.

### 2.5. Supplementation with linoleic acid abrogates the effect of LIX1 downregulation on mitochondrial morphology and ultrastructure in GIST cells and restores YAP1/TAZ and KIT levels

PHBs form large, hetero-oligomeric ring complexes composed of multiple PHB1 and PHB2 subunits [38]. PHB complexes might function as scaffolds in the IMM by regulating the lateral distribution of membrane lipids and proteins, thereby defining functional membrane domains [39]. Genetic interaction data linked the function of PHB complexes to cardiolipin, a dimeric phospholipid with four fatty acyl chains (mostly linoleic acid) and found predominately in the IMM. Cardiolipin levels are significantly reduced in mitochondria of cells lacking PHB2 [40], but they can be restored by linoleic acid supplementation in the growth medium [41]. As PHB2 levels were reduced in mitochondria of cells in which LIX1 was silenced (Fig. 5E), we added linoleic acid or vehicle (DMSO) in the medium of GIST-T1-*ShLIX1* and control cells for 12h. Analysis of the mitochondrial morphology and cristae organization showed that untreated (vehicle) GIST-T1-*ShLIX1* cells displayed an elongated mitochondrial network, whereas linoleic acid-supplemented GIST-T1-*ShLIX1* cells harbored more fragmented mitochondria, as observed in untreated GIST-T1-*Scrambled* cells (Fig. 6A). Supplementation with linoleic acid slightly increased the number of mitochondria in GIST-T1-*ShLIX1* cells, and decreased the mean elongation and interconnectivity scores (Fig. 6B). Next, analysis of the IMM architecture by TEM indicated that untreated GIST-T1-*ShLIX1* cells harbored vesicle-like cristae without junctions, whereas linoleic acid-supplemented GIST-T1-*ShLIX1* cells displayed lamellar cristae, like control cells (Fig. 6C). Thus, linoleic acid supplementation abolished the mitochondrial defects caused by LIX1 silencing and restored GIST cell identity, as indicated by the marked increase in TAZ and KIT expression and the decrease of αSMA (SMC marker) (Fig. 6D-F). This demonstrates that LIX1 controls GIST tumorigenesis by regulating mitochondrial functions.

**Figure 6.**
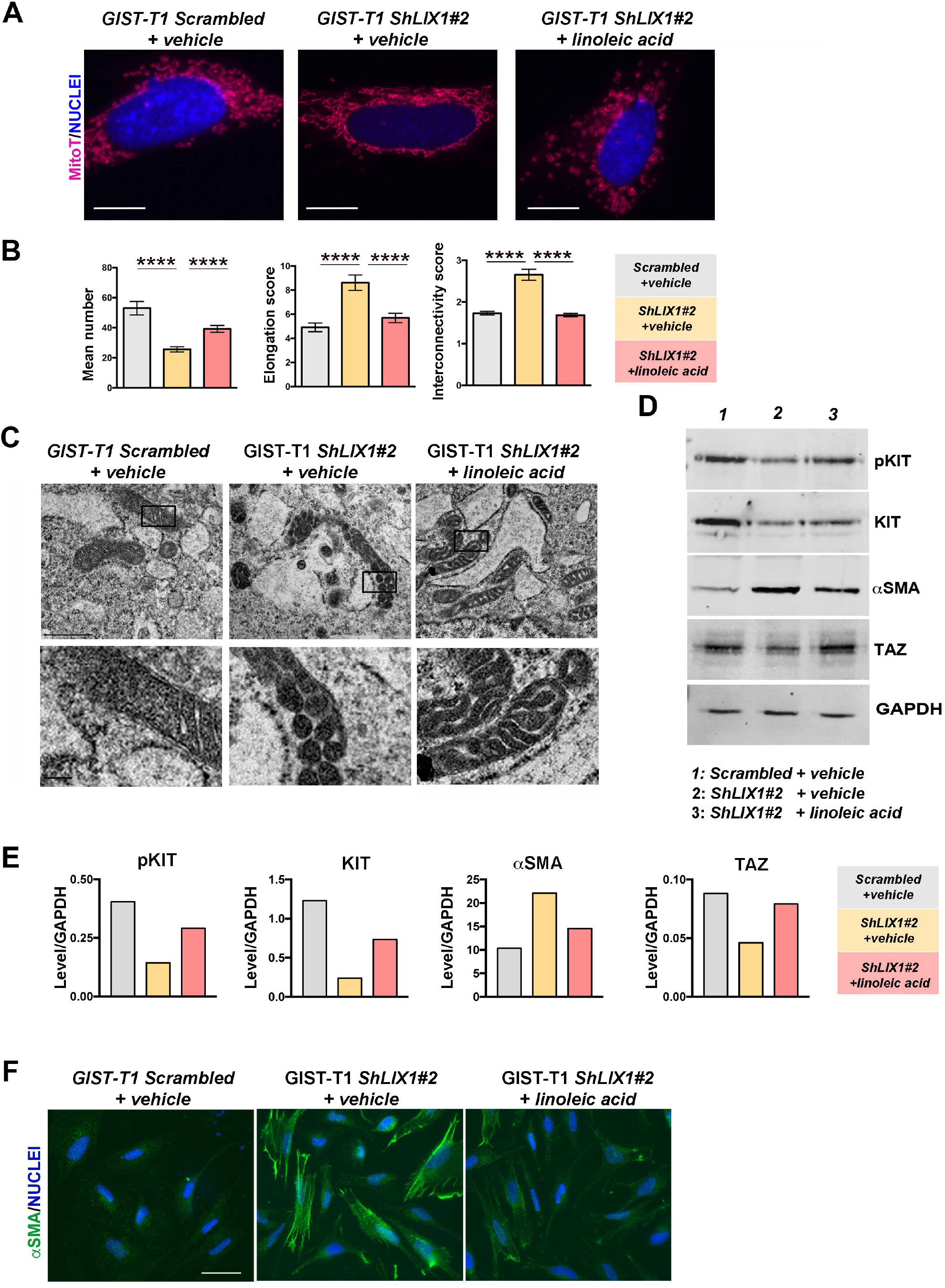
Incubation of GIST-T1 cells with linoleic acid abrogates the effect of LIX1 silencing on mitochondrial morphology and ultrastructure and restores YAP1/TAZ and KIT levels. (**A**) MitoTracker staining of GIST-T1-*Scrambled* (+DMSO vehicle), GIST-T1-*ShLIX1#2* (+DMSO vehicle), and -*ShLIX1#2* cells (+50μm of linoleic acid). Nuclei were visualized with Hoechst. Scale bars, 10μm. (**B**) Quantification of the MitoTracker data with the Mito-Morphology Macro in Image J. Values are the mean ± SEM of GIST-T1-*Scrambled* (+DMSO vehicle) *(n=25)*, GIST-T1-*ShLIX1#2* (+DMSO vehicle) *(n=28)*, and GIST-T1-*ShLIX1#2* cells (+50μm of linoleic acid) *(n=28)*; ****P <0.0001 (two-tailed Mann–Whitney test). (**C**) Ultrastructure of mitochondria in GIST-T1-*Scrambled* (+DMSO vehicle), GIST-T1-*ShLIX1#2* (+DMSO vehicle), and -*ShLIX1#2* (+50μm of linoleic acid) cells. Lower panels: magnification of the areas in the black boxes in the upper panels. (**D**) Representative western blot showing phosphorylated KIT (pKIT), KIT, αSMA and TAZ levels in GIST-T1-*Scrambled* (+DMSO vehicle), GIST-T1-*ShLIX1#2* (+DMSO vehicle) and -*ShLIX1#2* (+50μm of linoleic acid) cells. Equal loading was verified by GAPDH expression. (**E**) Quantification of the western blot bands to determine the expression levels of phosphorylated KIT (pKIT), KIT, αSMA and TAZ relative to GAPDH in the different experimental conditions. (**F**) Immunofluorescence analysis of αSMA expression in GIST-T1-*Scrambled* (+DMSO vehicle), GIST-T1-*ShLIX1#2* (+DMSO vehicle), and -*ShLIX1#2* (+50μm of linoleic acid) cells. Nuclei were visualized with Hoechst. Scale bars, 50 μm.

## 3. Discussion

We previously identified LIX1 as a master regulator of digestive mesenchymal progenitor specification through the control of the Hippo effectors YAP1/TAZ [12]. LIX1 is upregulated in GISTs and is a poor prognostic factor. Upon *LIX1* inactivation, GIST cells lose KIT expression, a marker of GIST cell identity, and they acquire features of differentiated SMCs. This is associated with reduced proliferative and invasive capacities. We also found that in GISTs, YAP1/TAZ signaling is implicated in KIT-mediated oncogenesis that is regulated by LIX1 [13]. However, the exact mechanisms by which LIX1 controls YAP1/TAZ levels and GIST development remained to be elucidated.

### 3.1. S-palmitoylation on cysteine 84 of LIX1 contributes to its anchorage to the outer membrane of mitochondria

We demonstrated a novel paradigm whereby LIX1-mitochondria tethering regulates mitochondrial ultrastructure and signaling pathways involved in GIST tumorigenesis. Indeed, LIX1 S-palmitoylation promotes its anchorage to the OMM, where it may be localized in detergent-resistant lipid domains. These raft-like micro-domains are enriched in the phospholipid cardiolipin that is present mainly in the IMM and at contact sites between the IMM and the OMM [42–47]. Interestingly, cysteine 84 that is required for LIX1 S-palmitoylation is conserved in all vertebrate LIX1 orthologs and in the Drosophila homologs, suggesting a conserved cellular regulatory mechanism modulating LIX1 functions.

### 3.2. LIX1 regulates the inner membrane architecture in GIST cells

In gastric mesenchymal progenitors and in GIST-T1 cells, LIX1 concentrates in mitochondria where it contributes to the regulation of their ultrastructure. Indeed, LIX1 silencing led to the generation of elongated mitochondria with altered cristae and reduced level of the GTPase OPA1, and also to a decrease in mitochondrial respiration and ROS production. As elongation and interconnectivity of mitochondria were slightly increased in deficient LIX1 cells, we hypothesized that this phenotype could be a consequence of mtROS decrease because ROS-mediated mechanisms have been involved in mitochondrial remodeling [48].

### 3.3. LIX1 controls the mitochondrial stability of PHB2

We found that LIX1 is required to control the mitochondrial stability of PHB2, a protein involved in cristae organization. Similar to LIX1 silencing, ablation of PHB2 destabilizes the long OPA1 isoform and disrupts cristae structure, a phenotype that led to decreased cell proliferation [35,49]. This effect on IMM ultrastructure, where the respiratory chain is localized and the activity of which is crucial for ATP and mtROS production to ensure uncontrolled proliferation of cancer cells [29–32], confers to LIX1 a pivotal role in GIST tumorigenesis.

Due to the sequence similarity with lipid raft-associated proteins [50–52], PHBs might function as IMM scaffolds to control the lateral distribution of proteins and lipids, and thereby define functional platforms [39,53]. The *de novo* synthesis of cardiolipin in mitochondria is followed by a remodeling process, in which cardiolipin undergoes cycles of deacylation and acylation mediated the TAFFAZIN protein. Remodeled cardiolipin contains predominantly linoleic acid, an unsaturated fatty acid. As cardiolipin and OPA1 levels are reduced in mitochondria lacking PHB2 [35,40], PHB2 might segregate cardiolipin into specialized membrane domains where it could bind to and stabilize proteins required for its remodeling [40] and to proteins involved in cristae organization (e.g. OPA1 and its regulatory protease OMA1) [40,49,54,55].

We found that addition of linoleic acid to GIST-T1 cells in which LIX1 was silenced normalizes mitochondrial morphology and ultrastructure. Therefore, LIX1 might act as a scaffold to control the stability of PHB2 and consequently cardiolipin distribution or remodeling at mitochondrial contact sites. More studies are required to precisely characterize the mechanisms by which LIX1 controls PHB2 stability in mitochondria.

### 3.4. Supplementation with unsaturated fatty acids counteracts the effects of LIX1 knock-down on mitochondrial ultrastructure and restores GIST cell identity

In GIST cells in which LIX1 was silenced, linoleic acid also restored YAP1/TAZ signaling and consequently KIT levels, indicating that LIX1-mediated mitochondrial remodeling controls YAP1/TAZ levels and subsequently GIST development/progression. In agreement, we found that modulation of the mitochondrial redox state is sufficient to promote KIT-positive GIST cells differentiation towards the SMC lineage. Moreover, previous studies showed that high mtROS levels activate signaling pathways that promote cancer cell proliferation [29–32], and YAP1 activity depends on mitochondrial morphology and ROS levels [56,57].

### 3.5. Conclusion

This study brings insights into how LIX1 controls GIST tumorigenesis. We found that LIX1 contributes to the maintenance of the cristae organization and mitochondrial function, which are essential for cancer cell proliferation. Therefore, the present study improves our understanding of LIX1 role in regulating cell dedifferentiation and plasticity in GIST pathophysiology, and opens new avenues for GIST treatment strategies. Moreover, our findings could be interesting also for other tumors, particularly neuroblastoma. This is a rare childhood cancer derived from neural crest precursors in which LIX1 expression is associated with bad prognosis and that results from autocrine activation of the SCF/c-KIT axis [58].

## 4. METHODS

### 4.1. Cell culture and Reagents

The GIST-T1 cell line was obtained from Cosmo Bio (Japan). This cell line was established from a metastatic human GIST that harbor a heterozygous deletion of 57 bases in exon 11 of KIT [59]. HeLa and HEK-293T cells were a kind gift of V. Baldin and A. Debant (CRBM, Montpellier France), respectively. Human gastric SMCs, provided by Innoprot Innovative (Spain), were grown at low density to maintain a synthetic undifferentiated phenotype. All cells were cultured in Dulbecco’s Modified Eagle medium (DMEM) supplemented with 10% fetal bovine serum and 1% penicillin/streptomycin. GIST-T1 stable cell lines (GIST-T1-*Scrambled* and GIST-T1-*ShLIX1#1* and GIST-T1-*ShLIX1#2*) were generated, as previously reported [13]. GIST-T1 stable cell lines were incubated with DMSO (vehicle) or with 50μM linoleic acid for 12 hours. All cell lines were routinely tested for the absence of mycoplasma contamination (VenorGeM OneStep Test, BioValley).

### 4.2. DNA plasmids and constructs

The LIX1 expressing plasmid was from Origene (#SC108332, USA). The cDNA encoding human LIX1 was subcloned in the pCS2HA plasmid to generate the HA-LIX1 fusion protein in which the HA-tag is at the N-terminus of LIX1. The pCS2HA-HA-LIX1 plasmid was used as template to generate the LIX1 variants in which cysteine 83 and 84 were substituted by serine residues with the QuikChange Site-Directed Mutagenesis Kit (Stratagene) according to the manufacturer’s protocol and the primers listed in supplemental Table S1.

### 4.3. Subcellular fractionation

Subcellular fractionation was performed as previously reported [60]. Briefly, cells were resuspended in Mitochondria Isolation Buffer (MIB, 200 mM sucrose, 10 mM Tris/MOPS, 1 mM EGTA/Tris, and protease inhibitors), and then lysed using a motor-driven homogenizer operating at 1,600 rpm. An aliquot of the resulting extract was used as whole cell lysate (Total). The remaining lysate was centrifuged at 600 g, 4°C for 10 min to remove any contaminants coming from the nuclear fraction. This supernatant (crude cytoplasmic fraction) was then centrifuged again at 7,000 g, 4° C, for 10 min to pellet mitochondria. The mitochondrial fraction was washed twice in MIB and resuspended in MIB. For the protease protection assay, purified mitochondria were resuspended in MIB and incubated or not with increasing concentrations of proteinase K (PK) on ice for 30 min. PMSF (2 mM) was then added to inhibit PK. For alkaline sodium carbonate extraction, 100 mg of mitochondrial fraction was re-suspended in 100 mM Na_2_CO_3_ (pH 11.5) and incubated on ice for 30 min. After centrifugation at 100,000g for 30 min, the supernatant (soluble proteins) and the pellet (membranes) fractions were harvested. Samples were then separated by SDS-PAGE followed by western blot analysis. For the detergent assay, purified mitochondria were resuspended in MIB and incubated or not with 1/10 of 4% digitonin, 2% NP40, 2% Triton X100, or 10% SDS. Samples were then centrifuged at 16,000g and the pellet (membranes) fractions were harvested. Samples were then separated by SDS-PAGE followed by western blot analysis.

### 4.4. Acyl-RAC assay

LIX1 S-palmitoylation was assessed using the acyl-resin assisted capture (Acyl-RAC) method and the CAPTUREome™ S-Palmitoylated Protein Kit (Badrilla, K010-310; Leeds, UK), as previously described [28]. Briefly, free thiol groups were first blocked. The remaining palmitate groups were then treated with a thioester cleavage reagent (cleaved fraction) or with a preservation reagent (control, preserved fraction). Then, proteins with newly released thiol groups were incubated with the capture resin. Both bound and unbound fractions were analyzed. For the experiment, 1 mg of total cell lysates was used. Samples were separated by SDS–PAGE followed by western blot analysis.

### 4.5. Cell fixation, immunofluorescence microscopies, quantification

Cells were seeded on fibronectin-coated (50 μg/ml per coverslips) coverslips, fixed with 4% paraformaldehyde in PBS containing 0.01% Triton X-100 for 10 min, blocked with 1% goat serum for 1h before incubation with primary (Table S2) and Alexa 350-, 488-, and 555-conjugated secondary antibodies (Life Science) in 0.1% goat serum. Nuclei were labeled with Hoechst (Invitrogen). Cells were imaged using a Zeiss AxioVision fluorescence microscope. Mitochondria were imaged using a Zeiss LSM780 confocal microscope. Live cell imaging was performed using a Leica SP8-UVconfocal microscope. For staining mitochondria, cells were incubated with 200 nM of MitoTracker™ Deep Red FM (Molecular Probes™) at 37°C for 30 min. Cells were then washed 3 times with DMEM, rinsed in PBS twice, and processed for immunofluorescence. Quantification was performed with a macro designed by Dagda et al. [34] in ImageJ. Mitochondrial superoxide ions were detected as described by Arena et al.[61]. Cells cultured on glass coverslips were incubated with DMEM and 5μM MitoSOX™ Red Mitochondrial Superoxide Indicator (Molecular Probes™) at 37°C for 10 min. Cell counting and pixel quantification were performed with ImageJ. Antibodies are listed in Table S2.

### 4.6. Seahorse analysis and high-resolution Oxygraph respirometry

Oxygen consumption rates (OCR) were measured using the XFe-96 Extracellular Flux Analyzer (Seahorse Bioscience). 0.5 ×10^5^ GIST cells (50-60% of confluence) were seeded on poly-D-lysine-coated XF96 plates in XF medium (non-buffered DMEM containing 10 mM glucose and 2 mM L-glutamine). Cells were left for 4-6h before starting the injection of drugs. OCRs were measured using the mitochondrial stress test in basal conditions and in response to oligomycin (1 μM), FCCP (1 μM), rotenone (100 nM) and antimycin A (1 μM; Sigma).

OCRs were measured also by high-resolution Oxygraph respirometry (Oroboros). The respiratory rates of 2-3×10^6^ cells were recorded at 37°C in 2 ml glass chambers. Cells were resuspended in respiratory buffer (0.5 mM EGTA, 3 mM MgCl_2_, 60 mM potassium lactobionate, 20 mM taurine, 10 mM KH_2_PO_4_, 20 mM HEPES, 110 mM sucrose, and 1 mg/mL BSA at pH 7.1) and permeabilized by incubation with digitonin (15 μg/10^6^ cells). Malate (5 mM) and pyruvate (5 mM) were added to provide NADH to complex I (CXI). The addition of succinate (10 mM) and rotenone (10 mM) allowed measuring respiration driven by complex II (CXII).

### 4.7. Transmission electron microscopy (TEM)

GIST-T1 cells were fixed in 2.5% glutaraldehyde in PHEM buffer, pH7.2, at room temperature for 1h, and post-fixed in 0.5% osmic acid in the dark and room temperature for 2h, dehydrated in a graded series of ethanol solutions (30–100%), and embedded in EmBed 812 using an Automated Microwave Tissue Processor for Electronic Microscopy (Leica EM AMW). Thin sections (70 nm; Leica-Reichert Ultracut E) collected at different levels of each block were counterstained with 1.5% uranyl acetate in 70% ethanol and lead citrate and observed using a Tecnai F20 transmission electron microscope at 200 KV at the CoMET MRI facilities, INM, Montpellier France. Three-dimensional reconstructions were performed with the AMIRA software.

### 4.8. Western blotting

Cells were resuspended in lysis buffer (20 mM Tris pH8, 50 mM NaCl, 1% NP40, cOmplete EDTA-free Protease Inhibitor Cocktail (Roche)) or in MIB (to analyze mitochondria). Protein lysates were boiled in SDS-PAGE sample buffer, separated by SDS-PAGE, and transferred to nitrocellulose membranes. Membranes were incubated with primary antibodies (Table S2).

### 4.9. Statistical analysis

Data were analyzed with two-tailed or one-tailed (when appropriate) Mann-Whitney tests and the GraphPad Prism 6 software. Results were considered significant when P <0.05 (*), P <0.01 (**), P <0.001 (***), or P <0.0001 (****).

## Declaration of Competing Interest

The authors declare that they have no competing interests.

## Author s’ Contribution

A.G. performed confocal imaging and biochemical analyses. C.A. performed High-Resolution Oxygraphy experiment and analysed the data; S.K. performed Seahorse experiment and analysed the data. C.C. performed electron microscopy experiments. S.F. and P.d.S.B. conceived the study. S.F., P.d.S.B., A.L., A.M., N.T. designed experiments with contributions from C.A., J.F., M.R., S.K. S.F. wrote the manuscript with inputs from A.G. and P.d.S.B. All authors read and approved the manuscript.

## Acknowledgements

This work was supported by the Association contre les Myopathies (AFM N°23800 to SF), by INSERM-Transfert (CoPoc N°MAT-PI-13315-A-02 to SF), and by Ligue Régionale Contre le Cancer Languedoc-Roussillon (2020 to P.d.SB). Work was also supported by institutional funds by the University of Montpellier, INSERM and CNRS. AG is a recipient of an AFM-Téléthon PhD studentship. We are grateful to members of Sandrine Faure and Pascal de Santa Barbara’s team for critical reading of the manuscript.

---

**Supplemental Figure S1.**
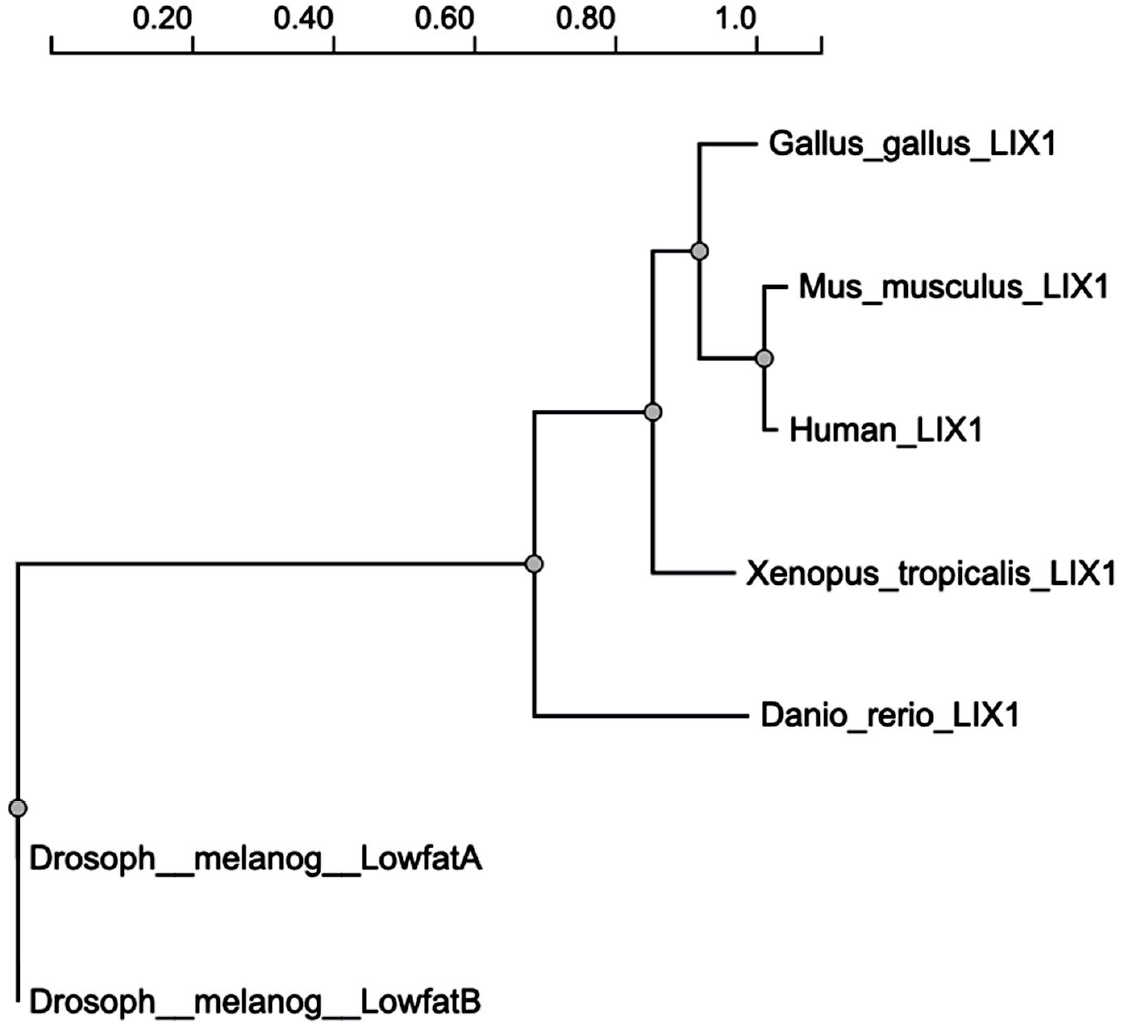
Phylogenetic tree using human LIX1 protein as query. At the end of each branch, the species names of the LIX1 orthologs are indicated. The accession number of each protein is indicated on the right. The scale bar represents the evolutionary distance of the tree branches (A.U.). The figure was generated with the MAFFT and FastME programs (https://ngphylogeny.fr/).

**Supplemental Figure S2.**
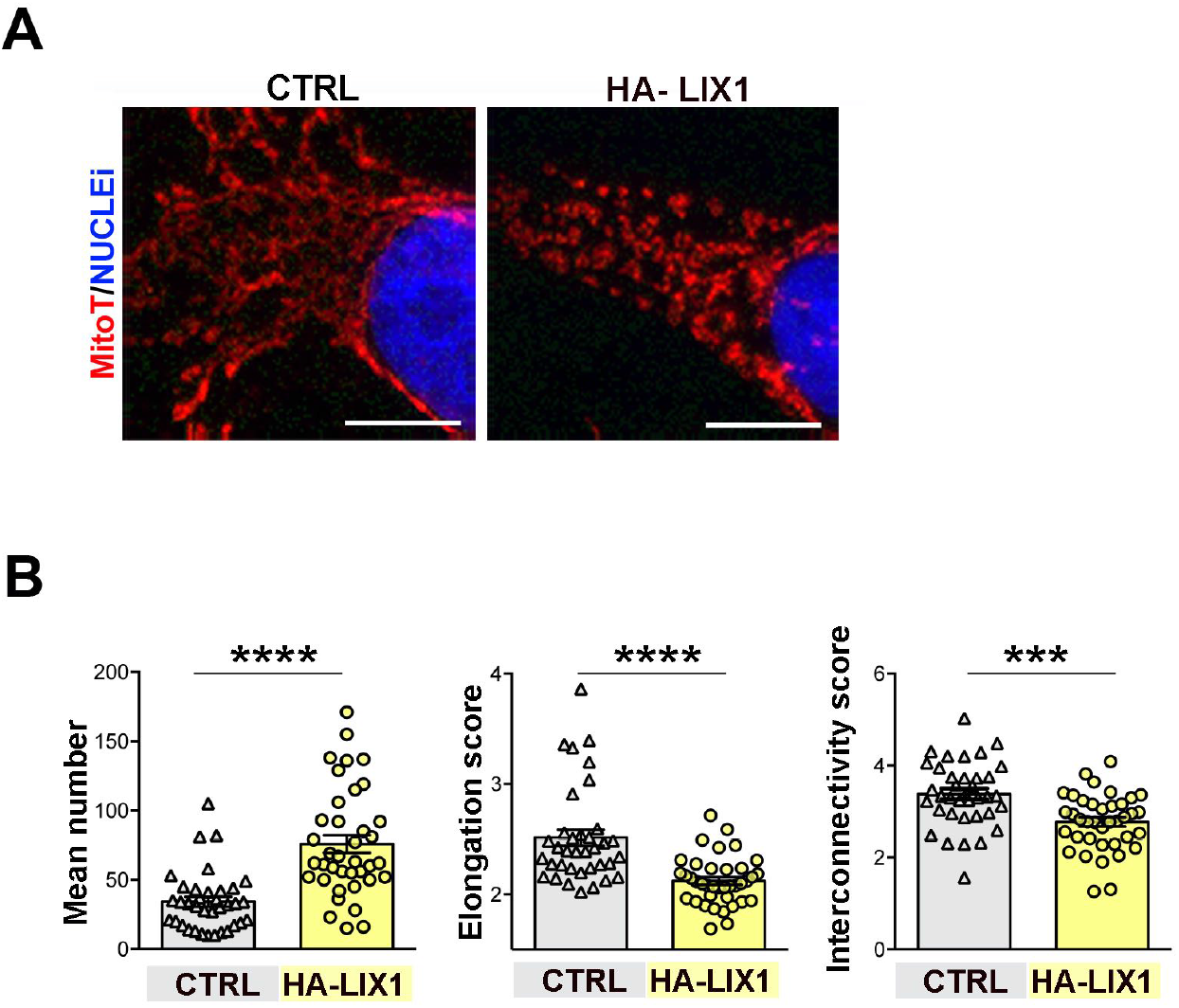
LIX1 promotes mitochondrial fragmentation of HeLa cells. **(A)** MitoTracker staining of control HeLa cells (CTRL) and HA-LIX1-expressing HeLa cells. Nuclei were visualized with Hoechst. Scale bars, 10μm. **(B)** Quantification of the MitoTracker data with the Mito-Morphology Macro in Image J. Values are the mean ± SEM of *n=36* HeLa control cells and *n=38* HA-LIX1-expressing HeLa cells. ***P <0.001; ****P <0.0001 (two-tailed Mann–Whitney test).

**Supplementary Figure S3.**
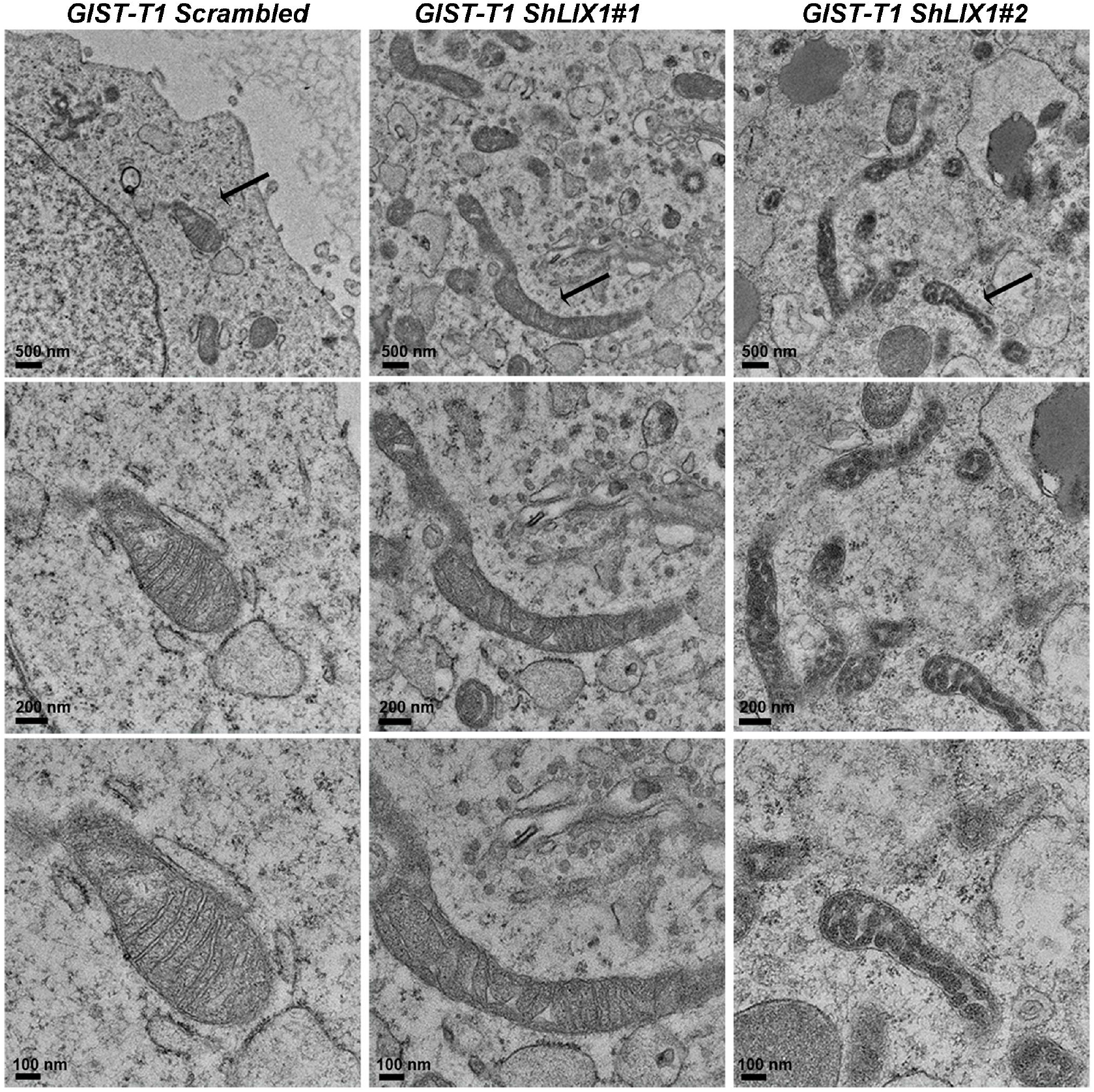
Ultrastructure of mitochondria in GIST-T1-*Scrambled*, GIST-T1-*ShLIX1#1* and -*ShLIX1#2* cells. These are tree other magnifications of the images shown in Fig. 5C. Scale bars are indicated on each panel. Black arrows point to the magnified areas.

## Notes

**Disclosure statement:** The authors disclose no potential conflicts of interest.

### Competing Interest Statement

The authors have declared no competing interest.

